# Informing grassland ecosystem modeling with in-situ and remote sensing observations

**DOI:** 10.1101/2024.06.28.601224

**Authors:** Johny Arteaga, Melannie D. Hartman, William J. Parton, Maosi Chen, Wei Gao

## Abstract

Historical grassland aboveground plant productivity (ANPP) was simulated by the DayCent-UV ecosystem model across the midwestern and western conterminous United States. For this study we developed a novel method for informing the DayCent-UV model and validating its plant productivity estimates for grasslands of the midwestern and western conterminous USA by utilizing a wide range of data sources at multiple scales, from field observations to remotely sensed satellite data. The model phenology was informed by the MODIS MCD12Q2 product, which showed good agreement with in-situ observations of growing season commencement and duration across different grassland ecosystems, and with observed historical trends. Model results from each simulated grid cell were compared to a remote-sensing ANPP modified version offered by the Analysis Rangeland Platform (RAP). This modified RAP ANPP calculation incorporated total annual precipitation, instead of mean annual temperature, as the control factor for the fraction of carbon allocated to roots. Strong temporal correlations were obtained between RAP and DayCent-UV, especially across the Great Plains. Good agreement was also found when the model results were compared with ANPP observations at the site and county level. The data produced by this study will serve as a valuable resource for validation or calibration of various models that aim to capture accurate productivity dynamics across diverse grassland ecosystems.

**Plain Language Summary:** This research used a computer model called DayCent-UV to simulate daily grassland growth across the central and western regions of the contiguous United States. To improve the agreement between the simulations and real-world conditions, we incorporated data from local field measurements and satellite imagery. This data helped determine the start and end dates of the growing season at each location. The simulated annual growth showed good agreement with satellite estimates from the Rangeland Analysis Platform (RAP), another computer application that monitors rangeland vegetation, and with local observations based on harvesting and weighing vegetation, particularly across the Great Plains. These results are valuable for validating and refining other computer models that aim to accurately simulate plant growth in grassland ecosystems; the predictions of these models are crucial for understanding the balance of carbon between plants, soils, and the atmosphere as the climate changes.

**Key Points:**

- The DayCent-UV model was used to simulate historical aboveground net primary productivity (ANPP) for different grassland ecosystems across the midwestern and western United States.
- MODIS MCD12Q2 was used to provide the phenology for the model.
- The Rangeland Analysis Platform (RAP) fraction of biomass production allocated to roots calculation was modified, resulting in a stronger agreement between its ANPP estimates and those from the DayCent-UV model.
- Site- and county-level ANPP observations were used to validate the model.

## 1 Introduction

Grasslands, defined as terrestrial ecosystems dominated by herbaceous and shrub vegetation and maintained by fire, grazing, drought and/or freezing temperatures, comprise ∼40 percent of the earth’s land surface excluding Greenland and Antarctica (White et al. 2000). In the United States, rangelands, similarly defined as lands on which the native vegetation is predominantly grasses, grass-like plants, forbs, or shrubs suitable for grazing or browsing use, comprise about 30% of the entire land cover, totaling about 770 million acres (https://www.nrcs.usda.gov).

Grasslands provide forage for domestic livestock and are habitat for wildlife. Additionally, grasslands store approximately 34 percent of the global stock of terrestrial carbon, with ∼90% of their carbon stored belowground as root biomass and soil organic carbon (White et al. 2000). Grasslands are also vulnerable to degradation from climate change impacts including more intense and frequent drought (Bardgett et al. 2021). Because grasslands are so vital to biodiversity, carbon sequestration, and human and animal welfare, it is important to quantify annual and long-term variation grassland plant productivity and its response to changes in climate.

Estimation of grassland plant productivity over a large region can be achieved using a combination of in-situ biomass measurements, eddy covariance flux tower data, remotely sensed satellite data, and ecosystem modeling. Field-based measurements of grassland productivity, such as those based on clipping vegetation at peak biomass, provide a direct quantification of above-ground net primary productivity (ANPP), but these measurements are labor intensive and limited in their spatial coverage. Flux tower data provide a reliable estimate daily net ecosystem exchange (NEE) (total C sequestration in plants and soils), but their spatial coverage across is not yet continuous and this data does not directly estimate net primary productivity (NPP) or ANPP. Phenocams are tower-mounted digital cameras that provide on-the-ground information about the phenology of rangeland plants. They can detect interannual variability in growing season commencement and duration and are important for designing management systems, but do not directly provide an estimate of NPP and are also limited in their spatial extent (Browning et al. 2019). Remotely sensed satellite data, such as NDVI, are frequent measurements (about every two weeks) that have existed for several decades at various resolutions and that detect long-term changes in the timing and magnitude of vegetation greenness over large spatial extents. This data has been used to estimate biomass (Morgan et al. 2016) and NPP (Hermance et al. 2015, Chen et al. 2019, Hartman et al. 2020, Jones et al. 2021). In particular, the Rangeland Analysis Platform (RAP) uses NDVI, a plant functional type cover dataset, and linear mixing theory to estimate herbaceous biomass in rangelands (Jones et. al, 2021). In turn, estimates of biomass derived from remotely sensed data are validated against on the ground measurements (Paruelo et al. 1997, Reeves et al. 2021).

Ecosystem models represent the co-evolution of both plants and soils and can tie together disparate data sources by using them as model input or for model calibration/validation. The models require regional sources of meterological data and soils data as inputs, phenological data (growing season start and end dates) as input or for model validation, and observations ANPP to compare to model predictions. Particularly, advanced models like the Community Land Model still struggle in defining the start and the end of the growing season, which is crucial to avoid bias in productivity (Li et al. 2022).

For this study we developed a novel method for informing the DayCent-UV model and validating its plant productivity estimates for grasslands of the midwestern and western conterminous USA by utilizing a wide range of data sources at multiple scales, from field observations to remotely sensed satellite data. We also demostrate how these myriad data sources are related. We used site-level biomass data at semi-arid and mesic grassland for detailed calibration of model parameters. We used MODIS satellite data with location-specific green up and green down dates to define growing season start and duration for DayCent-UV, and we show how this MODIS phenological data is consistent with PhenoCam and flux tower data. We also used other site-level ANPP data, along with county-level peak biomass data from the Natural Resource Conservation Service (NRCS), and RAP NPP estimates to validate the model predictions of ANPP at multiple spatial scales.

The goal of this research was to accurately simulate the interannual variability of grass plant productivity in the grassland regions of the midwestern and western conterminous USA in order improve our understanding of how these ecosystems respond to changing climatic conditions. Previous studies have shown that grassland productivity is strongly correlated to growing season precipitation or actual evapotranspiration (Sala et al. 1988, Chen et al 2019). Interannual variability in precipitation and production has been linked to longer cycles in sea surface temperatures (Chen et al. 2017). This research serves as a foundation for simulating climate change impacts on grassland ecosystems, as well as to support advances in long-term remote-sensing products aiming to assit land managers and to motivate a more exhaustive temporal and spatial validation in regional grasslands simulations.

## 2 Materials and Methods

### 2.1 Spatial domain

The spatial distribution of the simulation grid, with 30 km x 30 km spatial resolution, covers most of the grassland ecosystems in the USA, including the Great Plains (Short Grass Steppe, Mixed Grass Prairie and Tall Grass Prairie), cold and warm desert ecosystems (Great Basin, Colorado Plateau and Chihuahuan Desert), inter-mountain grasslands (Montana valley, Wyoming Steppe and Palouse), and the annual grasslands in California (Figure 1). Each of the 2712 grassland grid cells were defined based on the USGS’s 24-category land cover classification map, adapted to the Climate-Weather Research and Forescated model (He et al. 2022).

**Figure 1.**
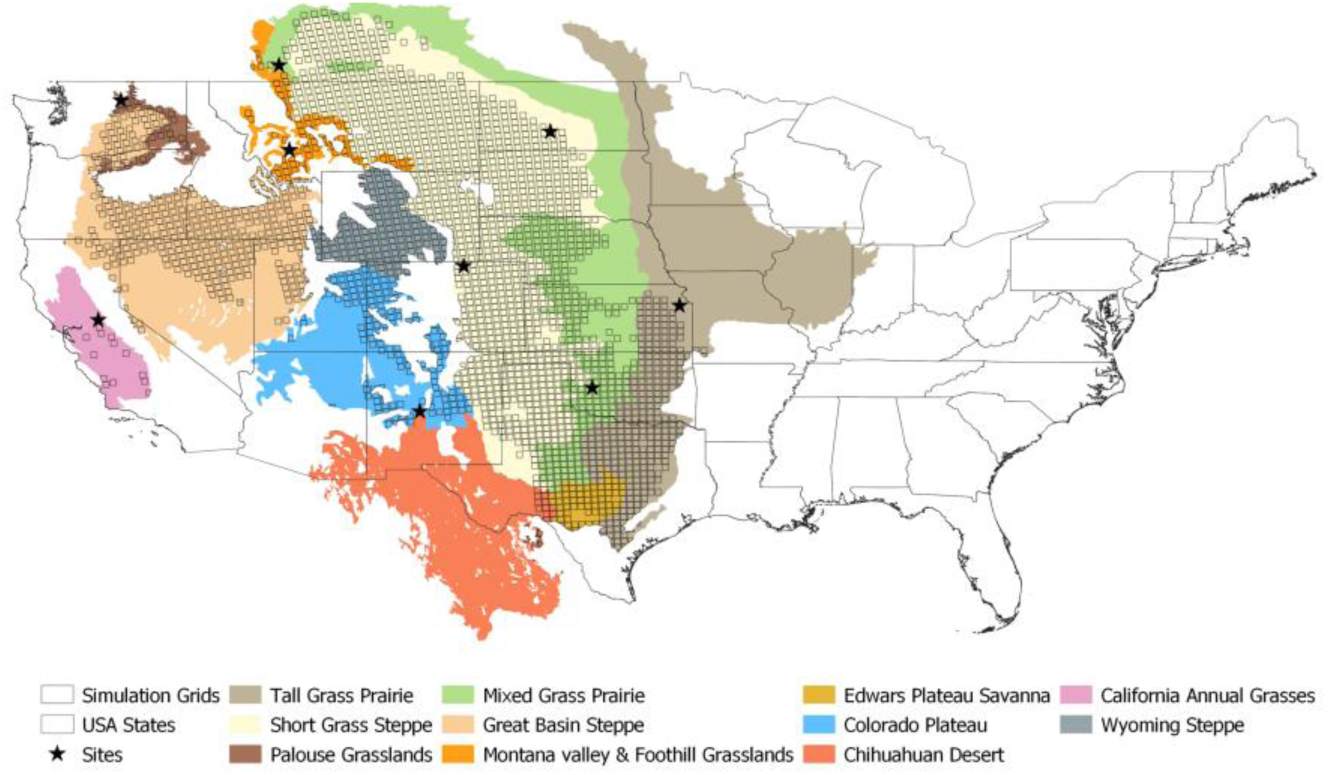
The grassland simulation grid (hollow black squares) overlaying different ecosystems in the western and midwest United States.The Great Plains region includes the Short Grass Steppe, Mixed Grass Priarie, and Tall Grass Prairie in the central part of the map from Canada (in the north) and Texas (in the south). The black stars indicate the locations where we validated e satellite-derived phenology with PhenoCam and other data. Map modified from Olson et al. (2001)

### 2.2 The DayCent-UV model

The DayCent ecosystem model is a process-based model that simulates daily fluxes of water, carbon and nutrients between the atmosphere, soil and vegetation (Parton et al. 1998). DayCent can simulate the plant production, soil carbon decomposition and formation, nutrient cycling, water fluxes, trace gas fluxes, and soil temperature dynamics across the vertical soil profile (Del Grosso et al. 2011), and is able to incorporate alterations introduced by the environment and management practices including fire, grazing, fertilizer application, tillage, and harvest.

In DayCent, the plant submodel computes the above and below ground biomass based on a plant’s maximum potential production, which depends on the solar incoming radiation, the light use efficiency of each plant type, water availability, and temperature. For grasses and other plant types, the total NPP corresponds to the maximum potential production reduced by nutrient limitation. The NPP is allocated to above-ground shoots and below-ground juvenilte and mature fine roots according to the shoot/root ratio, which is a function of soil moisture and nutrient availability (Parton et al. 1987, Gherardi & Sala 2020). In this work we used the DayCent-UV version, which also simulates the C losses associated with the photodegradation of surface and standing dead plant litter due to incident ultra-violet radiation (Chen et al. 2016). We used a point version of the model for calibrating parameters at the site-level and a gridded version for the regional simulations; these two versions differ only in the input data supplied to the models.

### 2.3 DayCent-UV optimization

To prepare DayCent-UV to simulate all grasslands in the region (Figure 1) we first optimized the point version of the model for two distinct grassland ecosystems: Central Plains Experimental Range (CPER) in the Shortgrass Steppe of Colorado and Konza Tallgrass Prairie in Kansas (Table 1) where long-term observations of aboveground plant productivity (ANPP) were available (Blair and Nippert 2024, Dorich et al. 2021). To minimize temporal bias and maximize the coefficient of determination (R²) between observations and model predictions of ANPP, we employed a Bayesian optimization approach to identify the optimal parameters for modeling annual grassland ANPP. For this optimization we used site-specific meterological inputs from the Applied Climate Information System (ACIS) Web Services (http://data.rcc-acis.org/), and specific soil properties data from STATSGO (Schawrz & Alexander, 1995). The observed annual average ANPP for three grazing treatments (light, moderate and heavy) were used for the CPER site while ANPP data from the annually burned ridge site were used for the Konza site.

**Table 1.**
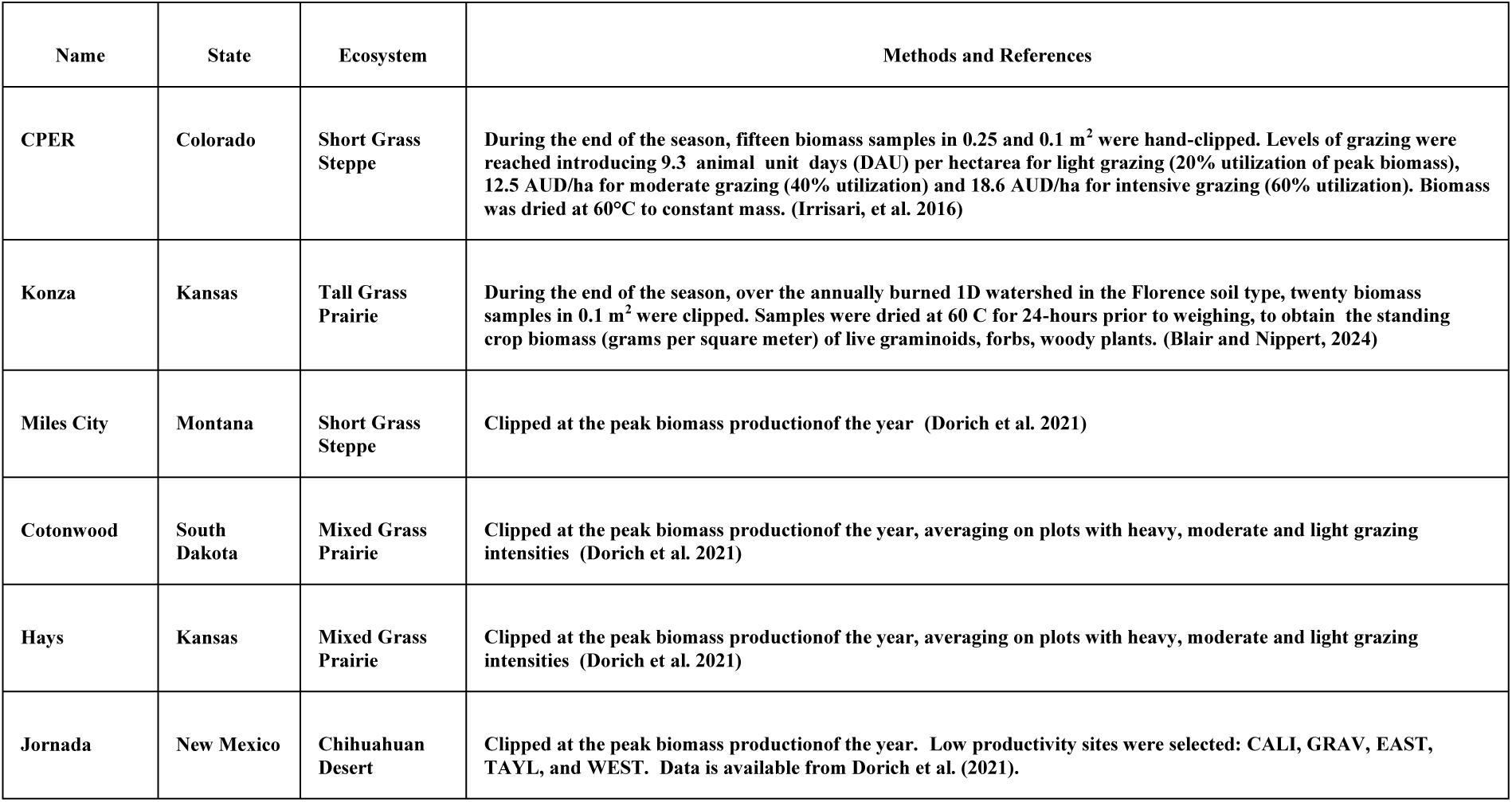
Sitelevel ANPP observations.

Detailed information on the necessary model adjustments to accurately simulate ANPP time series for both sites is provided in Section SP1 of the supplementary material. Live root biomass at both sites was assumed to be around 750 g biomass m^-2^, while the ratio of live root production to live shoot production was assumed to be 1.2 and 0.9 for CPER and Konza, respecitively (Milchunas & Lauenroth 2001, McCulley et al. 2009). In addition to calibrating plant growth parameters through the Bayesian optimaization, we adjusted live shoot death rates in the model so the simulated live leaf biomass for CPER and Konza followed the patterns of observed MODIS NDVI data at the two sites. Grassland plant attributes for each of the 2712 simulated grid cells, with the exception of annual grasslands in California, were assigned based on historical annual mean precipitation. Grid cells with annual precipitation below 500 mm were assigned plant attributes from the calibrated CPER site, while those exceeding 500 mm precipitation were assigned plant attributes from the Konza site. For the California annual grasslands we used plant parameters determined from previous work (Ryals et al. 2015).

### 2.4 DayCent-UV gridded inputs

Daily meteorological inputs from 1979-2015 for each grid cell in this study were obtained from the National Centers for Environmental Prediction North American Regional Reanalysis (NCEP, 2005). The daily metereological inputs necessary to run the gridded DayCent-UV model included precipitation, minimum air temperature, maximum air temperature, relative humidity, wind speed, and incoming solar radiation.

The vertical soil structure for each grid was based on the ten-layer configuration of the Common Land Model (Dai et al., 2003). Its hydraulic and textural properties were derived from a combination of two datasets: the Continental United States Multi-Layer Soil Characteristics Dataset (CONUS-SOIL) and the Food and Agriculture Organization of the United Nations Educational, Scientific, and Cultural Organization (FAO-UNESCO) Soil Map of the World (Liang et al., 2005).

### 2.4 Phenological Parameterization

We used the MODIS MCD12-Q2 phenology product to define the grid cell-specific growing season parameters used by the DayCent-UV model (Friedl et al. 2022). The ’GreenUp’ band, representing the first date when the EVI timeseries surpassed its 15% amplitude, was used to identify the growing season start. Similarly, the ’MidGreenDown’ band, signifying the last date when the EVI timeseries dipped below its 50% amplitude, determined the growing season’s end. MODIS MCD12-Q2 phenology product also provides the dates of peak biomass, Senescence, and Dormancy which we did not use. For each simulated grid, we extracted the mean and standard deviation for the Greenup and MidGreenDown dates over a 20-year period (2001-2020) from all MODIS pixels within the grid. The mean Greenup date minus one standard deviation determined the potential start of the growing season and the mean MidGreenDown date determined the onset of plant senescence. The actual start of the growing season was simulated on or after the potential start date and varied each year once air temperatures exceeded the threshold for growth.

To determine which of the MODIS MCD12-Q2 derived phenology values were most appropriate for our study, we selected nine sites across diverse grassland ecosystems (black stars in Figure 1). Daily green chromatic coordinate (GCC) data from the PhenoCam Dataset version 2.0 (Seyednasrollah et al. 2019) and daily Gross Primary Productivity (GPP) or Net Ecosystem Exchange (NEE) data from AmeriFlux eddy covariance towers were used for comparison. GPP reflects light use efficiency for carbon fixation, making it suitable for phenology assessment. The start of the growing season was defined as the date when the GPP timeseries began to rise from zero, while the end of the growing season was defined as its decline to zero. For NEE, the start of the growing season was defined when the NEE flux drops from positive to negative while the end of the growing season was defined when NEE becomes positive again.

To minimize the influence of non-grassland areas, the MODIS MCD12-Q2 products were masked using the annual grassland classification from the MODIS MCD12Q1 Version 6.1 product by the University of Maryland (UMD) (Friedl et al., 2022b). This process, performed through the Google Earth Engine platform, ensured the final processed time series represented only the relevant grassland areas.

### 2.5 ANPP estimated by the Rangeland Analysis Platform (RAP)

The Rangleland Analysis Platform (RAP) was developed to assist and monitor rangeland across USA (Jones et al. 2021). It estimates annual NPP by composing daily NDVI time series from 16-days Landsat imagery. RAP products have a 60m pixel resolution and a temporal window from 1986 to the current time. To estimate anuual ANPP, RAP multiplies the annual NPP by the fraction of carbon allocated in shoots 𝑓_𝐴𝑁𝑃𝑃_ , based on the Hui and Jackson empirical relationship (Hui & Jackson, 2006)

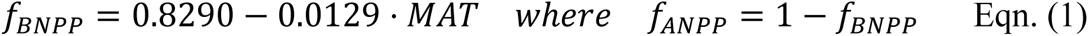

Here, 𝑓_𝐵𝑁𝑃𝑃_ is the fraction carbon allocated in roots and MAT is the mean annual temperature (°C). We propose an alternative approach incorporating the empirical relationship established by Gerardhi & Sala (2020). Their research suggests a stronger correlation between ANPP and annual precipitation (APPT, mm) compared to the formula in Eqn.(1), given by

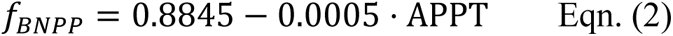

This relationship was also supported by an independent study in Sun et al. (2021).

We regridded RAP NPP estimates obtained from the RAP Google Earth Engine version (http://rangeland.ntsg.umt.edu/data/rap/rap-vegetation-cover/) to our 30 km x 30 km grassland grid and calculated two estimates of ANPP using Eqn. (1) and Eqn. (2). For Eqn. (2) we accessed the GRIDMET database to get the annual precipitation (Abatzoglou, 2013). We evaluated our proposed alternative RAP ANPP estimate and the original RAP ANPP calculation against DayCent-UV results and against site-level and county-level ANPP data.

### 2.6 Site and county level ANPP observations

We assessed the DayCent-UV model’s ability to simulate site-level temporal variability by comparing in-situ ANPP observations from long-term experiments (Table 1) with the corresponding modeled ANPP values from the nearest simulated grid cell. Note that although the point version of DayCent-UV was used to calibrate the plant model for CPER and Konza using local meterological and soils data, the grid cell-level results that we compared to in-situ observations for CPER, Konza, and the other four locations used grid cell averaged meteorological inputs and soil properties.

Annual ANPP in the period 2004 – 2014 were collected every few years by the Natural Resource Conservation Service (NRCS) across multiple sampling locations in each county in the Great Plains and were averaged by county to compute mean county-level ANPP (Chen et al. 2019). We averaged each simulated grid cell falling within each county to validate the DayCent-UV and RAP ANPP estimates against the NRCS data.

## 4 Results

### 4.1 Bayesian optimization

Given that seasonal precipitation is a major driver of annual ANPP variability (Lauenroth and Sala, 1992), we calibrated the DayCent model at long-term experimental sites representing contrasting precipitation regimes, CPER and Konza (Table 1). The best fit parameters for the CPER and Konza sites were obtained after performing a Bayesian optimization, constrained to a minimum bias and high temporal correlation (𝑅^2^) between site-level annual observed and simulated ANPP. The best temporal performance in both sites had 𝑅^2^ values close to 0.5 (Figure S1). Though simulated above ground shoot biomass was not directy used in the optimization process, continuous simulated above ground shoots coincided well with MODIS-NDVI values, particularly at CPER (Figure S1).

### 4.2 Evaluation of phenology data

Using an dependable source of data to define model phenology is crucial in regions where the start and the end of growing season varies spatially. We utilized the ’GreenUp’ and ’MidGreenDown’ bands from the MODIS MCD12-Q2 product to define the start and the end of the growing season, respectively.

We confirmed that the MODIS GreenUp and MidGreenDown dates were consistent with other available ecosystem data. To validate these key dates we showed that there was good agreement in the start/end of the growing season between MODIS phenology and the rise/fall in photosynthesis activity informed by GPP (green filled) or NEE (gray filled) (Figure 2). We observed a strong agreement between the temporal patterns in sites with GPP and GCC information. This agreement allows us to validate MODIS phenology dates at sites with only PhenoCam information. Overall, for the nine sites representing diverse grassland ecosystems, MODIS GreenUp and MidGreenDown dates successfully captured the start and end of the growing season, with the exception of sites across the central Great Plains (Oklahoma and Colorado). These central Great Plains sites exhibited greater inter-annual variability, particularly in the end-of-season detection.

**Figure 2.**
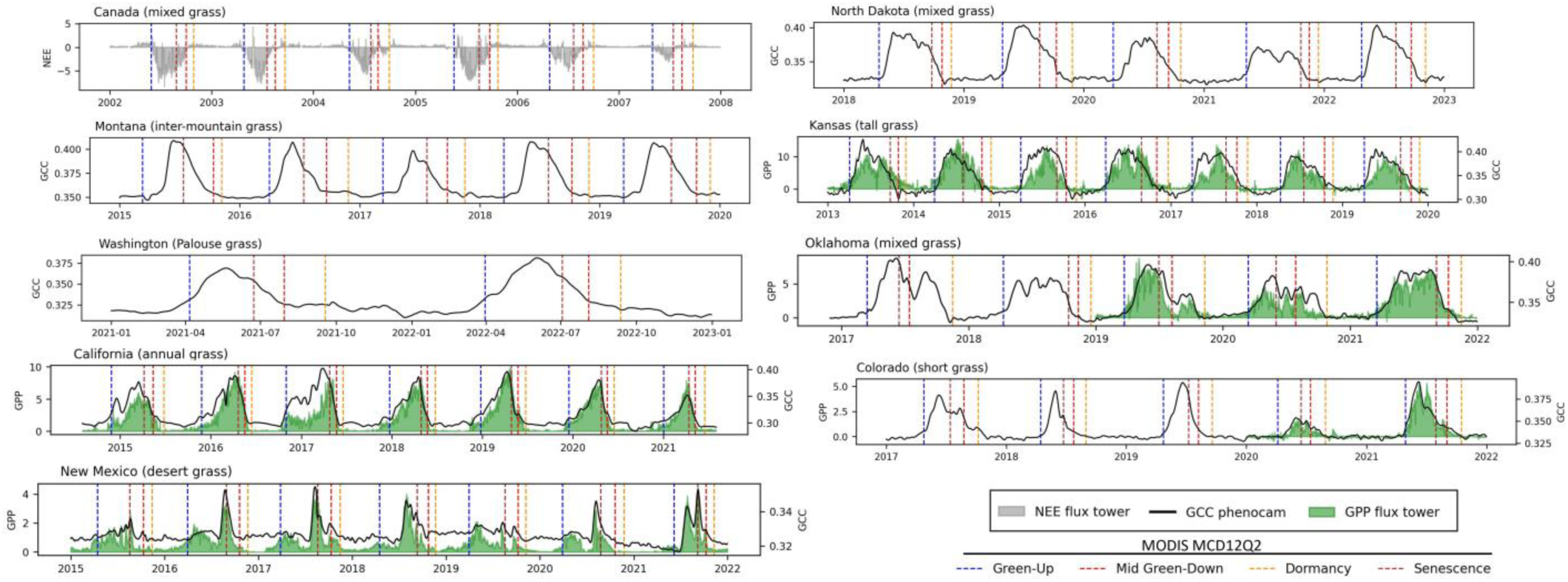
Validating MODIS MCD12-Q2 phenology across different ecosystem sites in Figure 1: Temporal variation for GPP in 𝜇𝑚𝑜𝑙/𝑚^2^𝑠 (green filled), NEE in 𝜇𝑚𝑜𝑙/𝑚^2^𝑠 (gray filled) and GCC (black solid line) and the corresponding MODIS dates indicating the Green-up (vertical blue dashed line), Senescence (vertical brown dashed line), Mid Green-Down (vertical red dashed line) and Dormancy (vertical orange dashed line).

We further investigated the utility of the Senescence and Dormancy bands from the MCD12-Q2 product. However, both bands exhibited limitations for our purposes, particularly in the Great Plains region. The Senescence band, defined when the EVI time series falls below 90% of its amplitude, typically occurred around one month earlier than the actual end of the growing season in the Great Plains (which usually falls around late August). This earlier timing leads to underestimation in plant productivity (results not shown). Conversely, the Dormancy band, defined when the -EVI time series drops below 15% of its amplitude, lags the actual end of the growing season by about a month, causing productivity overestimations (results not shown).

To explore the potential for shifts in the start and end of the growing season between 2001 and 2020, we performed a temporal trend analysis for the GreenUp and MidGreenDown dates over this 20-year period (Figure S2). The start of the growing season shows a negative trend in the northern Great Plains, indicating an earlier onset of the growing season by about one day per year. In addition, a positive trend, indicating a later onset of the growing season, was observed in the Mixed Grass Prairie in North Kansas. Regarding the end of the growing season, a delayed senescence was observed in South Dakota and North Dakota.

### 4.3 Comparing DayCent-UV simulated ANPP against RAP estimates

After optimizing plant growth parameters, and specifying the average start and end of the growing season for each grid cell, each grassland grid cell in the midwestern and western USA was simulated at a daily time scale from 1979 until 2015 using grid-cell specific weather and soil information. The simulated average annual ANPP and RAP remote-sensing estimations using Eqn. (2) to estimate ANPP from total NPP (1986-2015) showed similar pattern across all simulated ecosystems (Figure 3). Starting from the west, ANPP for California grasslands averaged from 150 to 300 g/m^2^ for both RAP and DayCent-UV. Simulated productivity for grasslands between California and the Great Plains was generally below 100 g/m^2^ and slightly higher than RAP estimations. Going from west to east across the Great Plains, the both RAP and DayCent-UV ANPP gradually increased from 100 g/m^2^, in the Short Grass Steppe, to values above 400 g/m^2^ in the Tall Grass Prairie.

**Figure 3.**
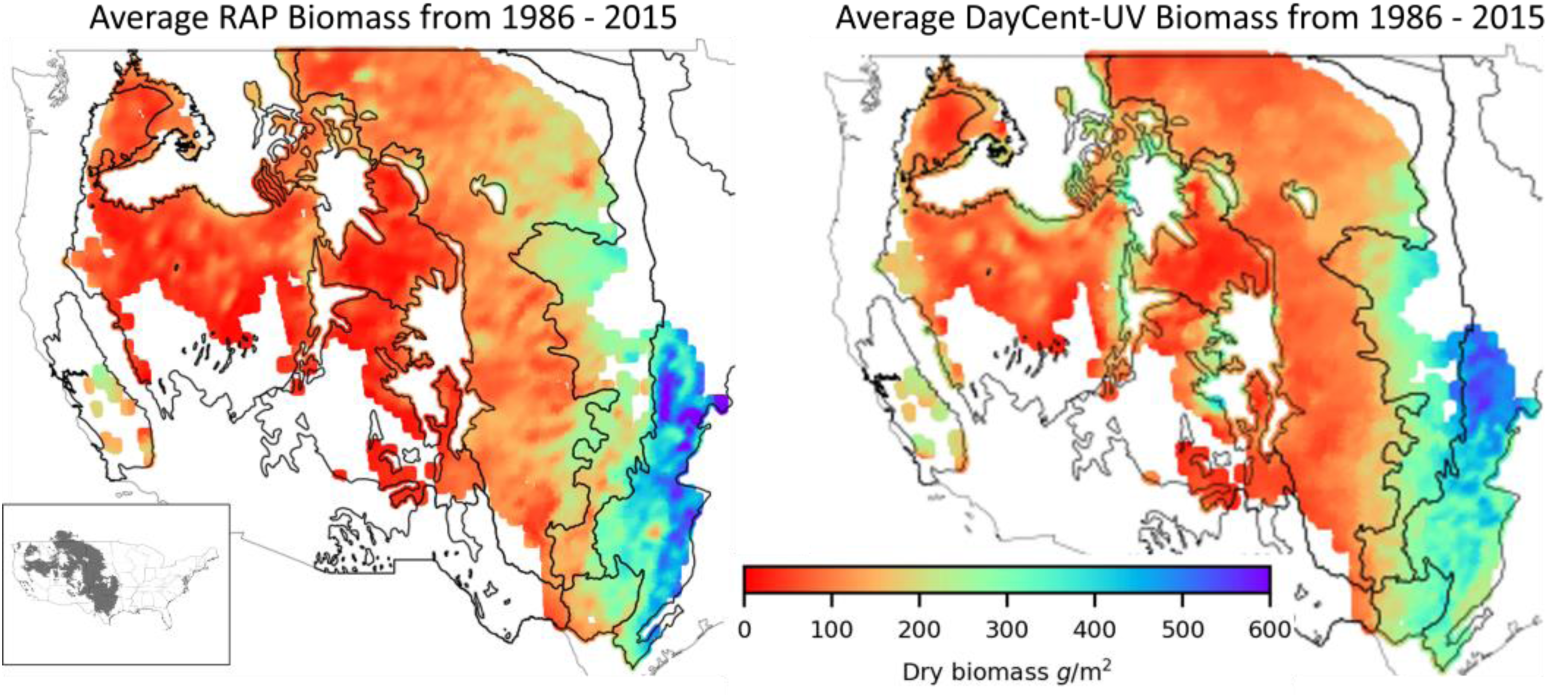
Average ANPP estimated by RAP (left) and by DayCent-UV model (right)

To evaluate the temporal performance of DayCent-UV model against RAP estimations, we calculated the coefficient of determination (R^2^) and relative bias (RB) for each simulated grid (Figure 4). The simulated ANPP for California grasslands showed a low RB, but a low temporal correlation (R^2^ < 0.3). For most of the grasslands in between California and the Great Plains, DayCent-UV’s ANPP was more than 100% of RAP’s, but we also found a high heterogeneity in the temporal correlation. We observed the best model agreement between DayCent-UV and RAP in the Great Plains, where DayCent-UV’s ANPP estimates were only around 30% less than RAP’s for most of the Short Grass Steppe (western Great Plains) and Tall Grass Prairie (eastern Great Plains), while DayCent’s APP estimates were higher than RAPS’ in most of Mixed Grass Prairie.

**Figure 4.**
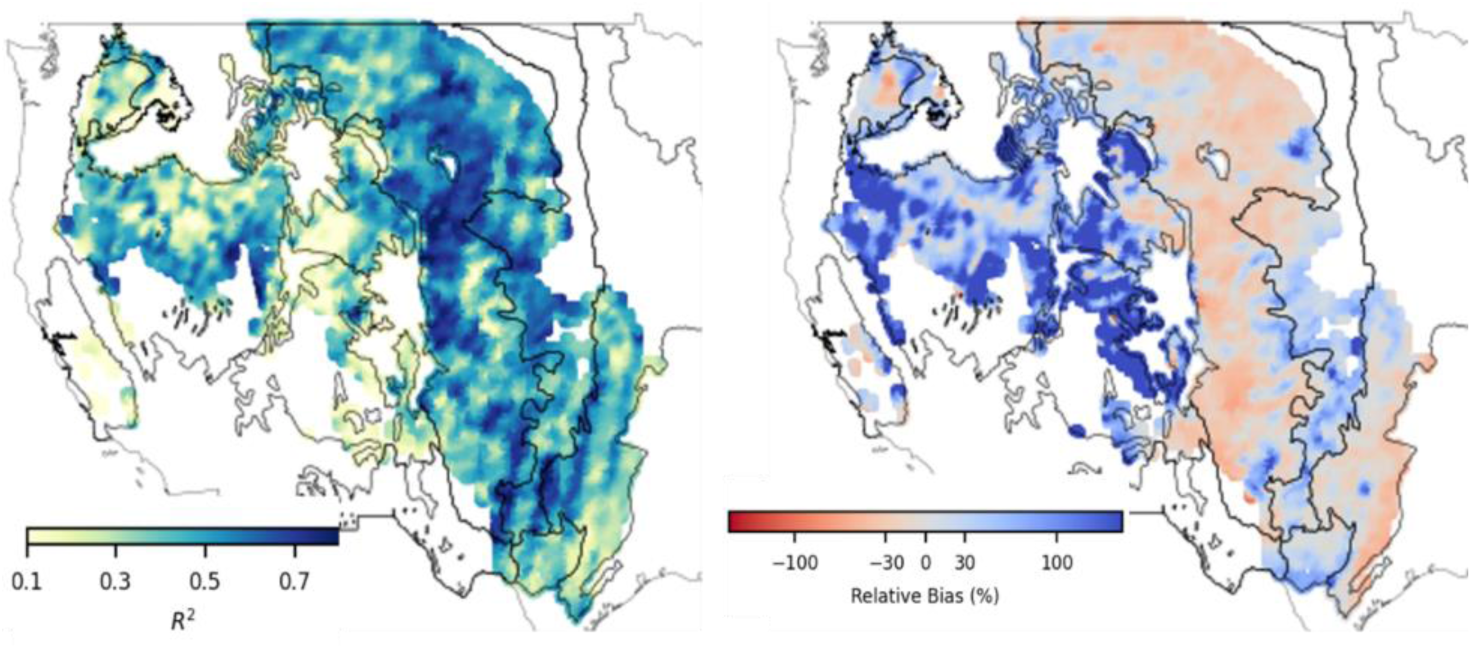
Temporal R^2^ between DayCent-UV ANPP and RAP ANPP (left) when Eqn 2 was used to estimate RAP ANPP. The corresponding relative bias between DayCent-UV ANPP and RAP ANPP (right)

The strength of the correlation between DayCent-UV and RAP was generally lower when RAP ANPP was estimated using the empricial relationship from Hui and Jackson (Eqn. 1) (Figure S3) than when using the one from Gerardhi and Sala (Eqn. 2) (Figure 4). In general, the grassland sites west of the Great Plains showed more regions with lower R^2^ in Figure S3 compared to Figure 4. For the Short Grass Steppe, Figure S3 results are similar to those of Figure 4 but with lower R^2^ at southern regions. For the Mixed and Tall Grass Prairie R^2^ values were lower and relative bias was increased in Figure S3 relative to Figure 4.

### 4.4 Validating county-level simulated ANPP against NRCS and RAP estimates

Maps of the county-level DayCent-UV simulated mean ANPP, mean RAP ANPP, and observed mean NRCS ANPP (2004 – 2014) for all of the Great Plains (Figure 5) showed a similar pattern with mean ANPP increasing from west to east across the Great Plains. The regression of the mean county-level ANPP between NRCS data and DayCent-UV predictions showed a good spatial correlation (R^2^ = 0.53). Mean DayCent-UV ANPP also had a high spatial correlation with mean RAP ANPP for these counties (R^2^ = 0.64). DayCent-UV ANPP estimates are most similar to NRCS ANPP for the Mixed Grass Prairie and Tall Grass Prairie in the eastern Great Plains.

**Figure 5.**
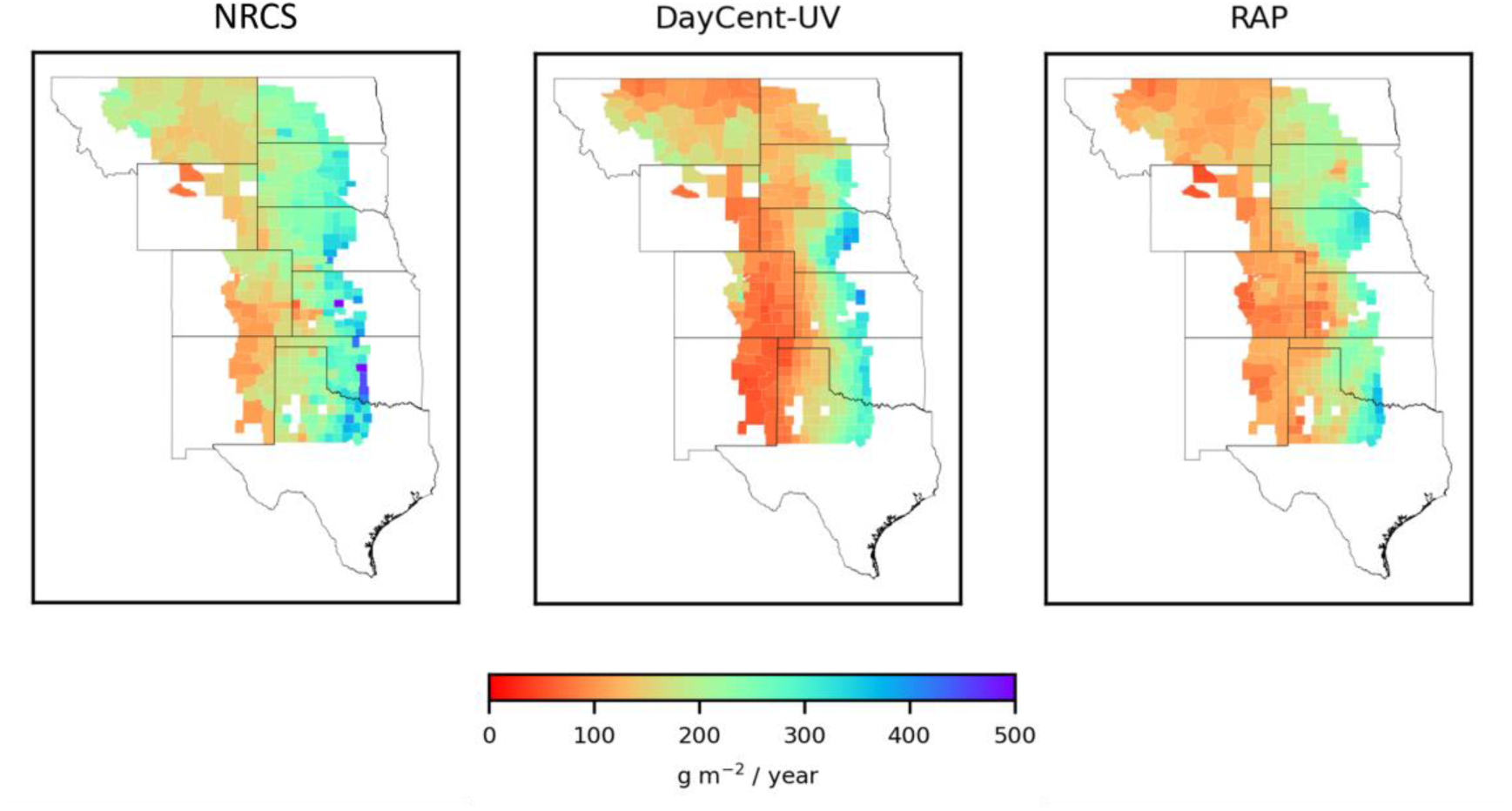
Average county-level ANPP for the period 2004 to 2014: NRCS data (left panel), DayCent-UV (middle panel), RAP ANPP (right panel).

However, in western counties, the NRCS ANPP shows higher productivity estimates compared to the DayCent-UV model. This bias is confirmed by the correlation analysis in counties with ANPP lower than 200 g/m² (Figure S4), where the slope of the trend line indicates an almost 60% greater ANPP by NRCS compared to DayCent-UV model. This may be due in part to the fact that NRCS data was not collected continuously each year (2004-2014), but DayCent-UV and RAP ANPP means include all years during this time period. In fact, DayCent-UV ANPP was more similar to RAP ANPP than it was to NRCS ANPP for this western region of the Great Plains.

### 4.5 Evaluating grid cell ANPP results against site-level data

We compared annual ANPP estimates from gridded DayCent-UV and gridded RAP against in situ observations at six sites in the Great Plains (Table 1) (Figure 6). Although DayCent-UV was calibrated for CPER and Konza using local meteorological drivers and soil texture, when DayCent-UV was run for the grid, it used grid cell-level weather drivers and soils data that were not completely consistent with site-level data. Nevertheless, DayCent’s grid-cell level estimates of ANPP for CPER had little bias compared to observations and were strongly temporally correlated to ANPP observations from 1991-2015. For Konza, DayCent-UV results were much higher than observed ANPP but were close to RAP estimates; however, grid cell-level sand content for the grid cell containing Konza was 10% while site-level sand content was 25% and higher sand content generally results in lower observed and simulated plant productivity. For Miles City, MT, DayCent-UV overestimated ANPP and showed the lowest R^2^ of the six sites, while RAP ANPP followed observations closely. The second lowest agreement between DayCent-UV and observations was for Jornada where both DayCent-UV and RAP underestimated observed ANPP; however the nearest grid cell to Jornada was 150 km away. For Cottonwood, SD, DayCent-UV underestimated observed ANPP. For Hays, KS, both DayCent-UV and RAP underestimated observed ANPP. Overall, DayCent-UV’s grid cell-level ANPP estimates seemed to correlate temporally with observed -site-level ANPP, showing similarities in interannual variability despite differences in scale, precipitation inputs, and soil texture, and with no consistent direction in the bias when it existed. DayCent-UV’s gridded ANPP estimates were often closer to RAPs grid cell level estimates than to site-level observations.

**Figure 6.**
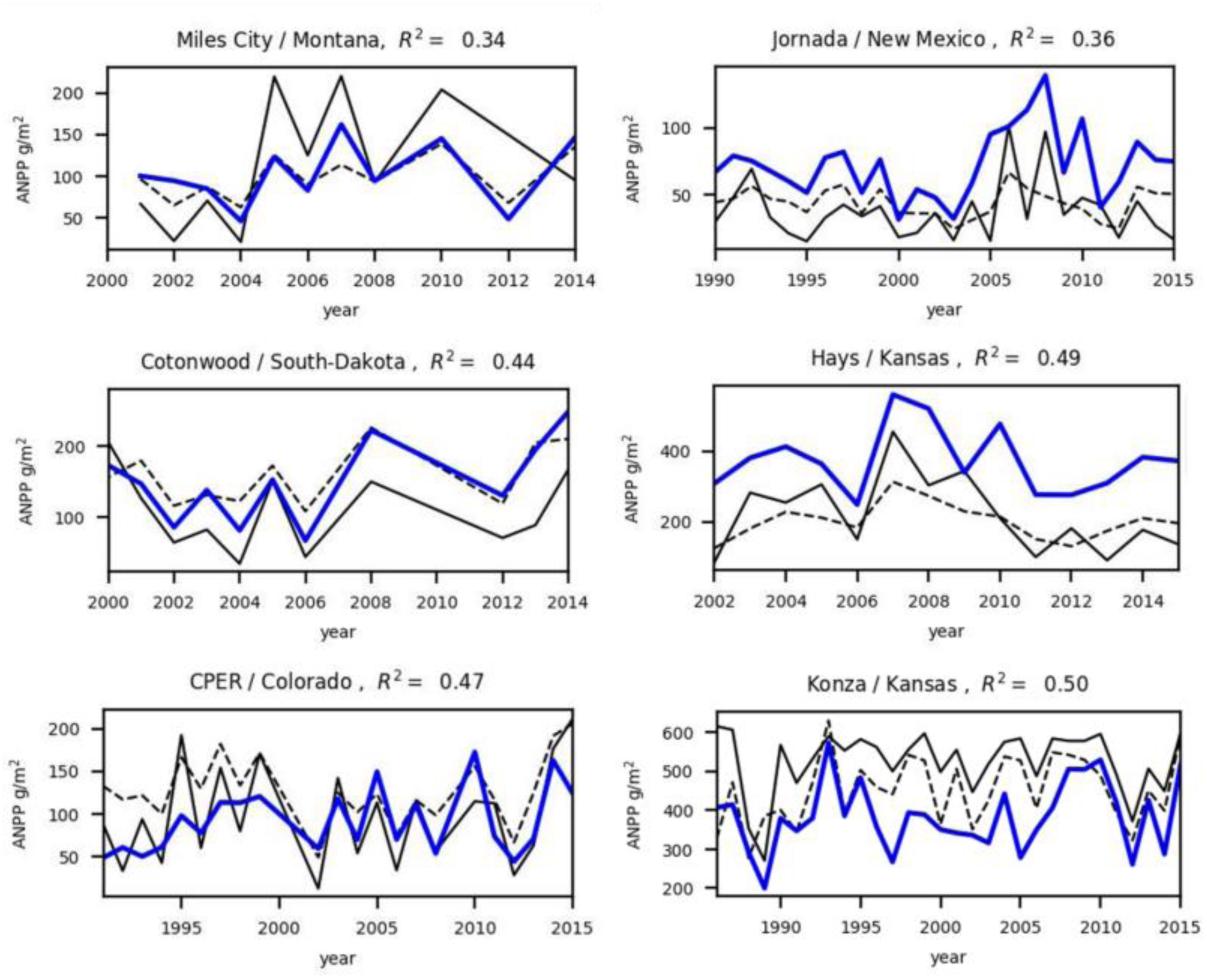
Site level comparison between ANPP field observations (blue lines) and the simulated ANPP (solid black lines) and RAP (dashed black lines) at nearest simulated grid. Temporal R^2^ are between site-level observations and DayCent-UV

## 5 Discussion

The DayCent-UV model was calibrated to simulate biomass productivity across various grassland ecosystems in the western and midwestern regions of the contiguous United States. We implemented a novel approach to define the growing season period within the model. This approach leverages phenological information derived from the MODIS MCD12-Q2 remote sensing product, specifically the Green-Up and Mid-Green-Down bands. The validity of this approach was supported by the observed -visual agreement between plant photosynthetic activity site records (GPP and GCC) and the growing season dates determined using the Green-Up and Mid-Green-Down bands (Figure 2). These findings align with those reported by Cui et al. (2019), who demonstrated good temporal correlations between GCC and daily MODIS time series data at four grassland sites.

Within the model, we defined the grid cell specific end of the growing season (beginning of the month-long senescence) as a fixed day. This day corresponds to the 20-year average of the Mid-Green Down date derived from the MODIS MCD12-Q2 product. After the simulation reached this predetermined day, aboveground live shoot biomass stoppped growing, gradually died, and was reduced to zero after one month. We found that the MidGreen down date was more appropriate for defining the onset of senescence for the DayCent-UV model than either the MODIS “Senescence” date or the MODIS “Dormancy” date. Not only did the MidGreenDown dates correspond to the end of the growing season as determined by the other ecological data (GPP, NEE, and phenocam GCC) (Figure 2), but DayCent’s estimates of plant productivity for the Great Plains (results not shown) were underestimated when the model used the “Senescence” date and were overestimated when the model used the “Dormancy” date to define the end of the growing season.

Recent papers suggest that potential impact of climate change in the Great Plains include increases in precipitation variability, earlier start to the growing season due to higher spring air temperatures, and increases in growing season air temperatures (Petrie et al. 2016, Post et al. 2022, Hayek & Knapp 2022, Burke et al. 2024). Chen et al. (2017) showed that variability of grassland plant production and growing season precipitation have increased from 1950 to 2020 for grasslands in eastern Colorado. In this paper, we utilized the observed remote sensing changes in plant phenology patterns (2001 to 2020) to inform the DayCent-UV model of the location-specific mean growing season onset and cessation. Our method defined the potential start of the growing season as the GreenUp date minus one standard deviation, allowing air temperature to trigger the actual start. However, the start of senescence was the same from year to year. This method may have been adequate for the timeframe we modeled (1979-2015) but trends in the start and end of the growing season will be increasingly important to include when simulating long-term changes in ANPP with climate change.

A trend analysis of the growing season onset (2001-2020) aligns with previous findings (Liu & Zhang, 2020) of earlier green-up in the northwestern Great Plains and a later spring onset north of Kansas (Figure S2). This earlier green-up in the north is likely attributed to a broad spring temperature shift across the region (Liu & Zhang, 2020). However, only a small portion of these sites exhibited a statistically significant trend (p < 0.05) towards an earlier start to the growing season, suggesting the site-specific dependence in phenology shifts (Post et al., 2022). The observed later spring onset north of Kansas could be associated with shifts in spring precipitation patterns (Ren et al., 2018). Furthermore, we observed a significant trend towards a later end of the growing season in the northern Great Plains (Figure S2). This is likely attributable to increased autumn temperature and precipitation (Ren et al., 2018).

During the model validation process, we evaluated the utility of the remote-sensing ANPP product offered by RAP. Our analysis revealed that incorporating an annual precipitation dependence into the fraction of carbon allocated to roots (Eqn. 2) improved RAP ANPP estimates compared to using a temperature dependence (Eqn. 1). With this improvement, DayCent-UV modeled productivity exhibited strong temporal correlation and low bias with RAP ANPP data in Great Plains grassland ecosystems, which aligns with the ecosystems used for model calibration (Figure 4). Model results for other grasslands, such as those in the Great Basin, Wyoming Steppe, Palouse Steppe, and Montana Valley, displayed a mixed pattern. In some locations DayCent-UV ANPP good temporal correlation and low bias when compared to RAP ANPP, while others exhibited higher bias. This finding presents a valuable opportunity to focus on long-term estimations in these understudied but important ecosystems. On the other hand, DayCent-UV ANPP for annual grasslands in California exhibited low bias but low temporal correlation with RAP ANPP. This discrepancy could be attributed to the way we accumulated productivity within each calendar year, since California grasslands start their growing season at the end of previous year. The high bias in DayCent-ANPP for cold and warm desert grasslands may be due in part to the limitations of remote sensing in arid areas, including the increasing concentration of invasive species in the Great Basin since 1990 (Smith et al., 2022), high co-dominance of shrubs and grasses in the Chihuahuan Desert (Smith et al., 2019), and the strong response of green-up and senescence to precipitation patterns in these ecosystems (Currier & Sala, 2022).

The gridded model productivity was also compared site-level ANPP observations (Figure 6) and average county ANPP observations across the Great Plains (Figure 5). While most of the site-level exhibited a good termporal correlation (R^2^ > 0.4), a high bias was observed. This bias is likely attributable to a combination of factors, including differences in precipitation between modeling grids and actual measurement sites, variation in soil layer properties across the study area, and the high dependence of plant productivity on topographic gradients (Nippert et al. 2011, Hoover et al. 2021). Regarding site-level measurements in desert grasslands, the underestimation of ANPP (Aboveground Net Primary Productivity) by both the model and RAP at the Jornada site in the Chihuahuan Desert (Figure 6) could be linked to the high spatial variability of ANPP observed in desert grasslands, as illustrated in Figure 2 by Muldavin et al. (2008). This evidence highlights the need for long-term simulations in this critical ecosystem to better quantify the accuracy of high-quality remote sensing products like RAP in capturing this high spatial variability (Smith et al., 2019).

The simulated DayCent-UV and the RAP ANPP estimates are intended to serve as a valuable tool for validation or calibration of various models that aim to capture accurate productivity dynamics across diverse grassland ecosystems. Examples includes models that uses machine learning for ANPP estimation (Sun et al. 2021 and Wiley et al. 2016), Land Surface Models performance against ANPP estimations derived from vegetation optical depth (VOD) (Fawcett et al. 2022), data assimilitation approaches in the Community Land Model (Fox et al. 2018), and process-based model driven by remote sensing observations like the Rangeland Carbon Tracking and Monitoring (Xia et al. 2024).

## 6 Conclusions

This study presented a long-term simulation of grassland productivity across the western and midwestern United States using the DayCent-UV model. The challenge to define the site phenology across such a broad region was addressed by leveraging the MODIS MCD12-Q2 product. The model exhibited strong temporal performance in most ecosystems compared to a newly proposed modification of the RAP ANPP estimate. This was particularly evident across the Great Plains and in specific regions like the Great Basin, Palouse Steppe, Montana Valley, and Wyoming Steppe. The simulated grid-level productivity was further evaluated against available site-level and county-level ANPP observations. This comparison revealed good temporal and spatial correlation, although some bias was observed at the site-level likely due to specific landscape features. The public availability of both the simulated dataset and the improved RAP ANPP estimate presents a valuable resource for supporting rangeland management practices and validating, calibrating, and performing data assimilation within various ecosystem models that simulate these important ecosystems. These model and validation improvements will be crucial for future assessments of climate change impacts on grassland ecosystems.

## Acknowledgments

This study is supported by the U.S. Department of Agriculture (USDA) UV-B Monitoring and Research Program, Colorado State University, under USDA National Institute of Food and Agriculture Grant 2022-34263-38472. This research is also partially supported by funds from USDA Grass-Cast Award 58-3012-0-021.

## Open Research

The grasslands grid cell distribution, the daily and annual DayCent-UV simulated ANPP, along with the ANPP estimates from the RAP product, are publicly available on Zenodo (https://zenodo.org/records/11165665)

## References

Abatzoglou, J. T. (2013). Development of gridded surface meteorological data for ecological applications and modelling. International Journal of Climatology, 33(1), 121–131.

Bardgett, R. D., Bullock, J. M., Lavorel, S., Manning, P., Schaffner, U., Ostle, N., … & Shi, H. (2021). Combatting global grassland degradation. Nature Reviews Earth & Environment, 2(10), 720–735.

Blair, J. and J. Nippert. (2024). PAB01 Aboveground net primary productivity of tallgrass prairie based on accumulated plant biomass on core LTER watersheds (001d, 004b, 020b) ver 17. Environmental Data Initiative.

Browning, D. M., Snyder, K. A., & Herrick, J. E. (2019). Plant phenology: taking the pulse of rangelands. Rangelands, 41(3), 129–134.

Burke, I.C., Yonker, C.M., Parton, W.J., Cole, C.V., Flach, K. and Schimel, D.S. (1989), Texture, Climate, and Cultivation Effects on Soil Organic Matter Content in U.S. Grassland Soils. Soil Science Society of America Journal, 53: 800–805.

Burke, M. W., Rundquist, B. C., & Caparó Bellido, A. (2024). Modelling vegetation phenology at six field stations within the US Great Plains: constructing a 38-year timeseries of GCC, VCI, NDVI, and EVI2 using PhenoCam imagery and DAYMET meteorological records. Theoretical and Applied Climatology, 1–17.

Chen, M., Parton, W. J., Adair, E. C., Asao, S., Hartman, M. D., & Gao, W. (2016). Simulation of the effects of photodecay on long-term litter decay using DayCent. Ecosphere, 7(12), e01631.

Chen, M., W.J. Parton, S.J. Del Grosso, M.D. Hartman, K.A. Day, C.J. Tucker, J.D. Derner, A.K. Knapp, W.K. Smith, D.S. Ojima, and W. Gao. (2017). The Signature of Sea Surface Temperature Anomalies on the Dynamics of Semi-arid Grassland Productivity. Ecosphere 8(12):e02069. 10.1002/ecs2.2069.

Chen, M., Parton, W. J., Hartman, M. D., Del Grosso, S. J., Smith, W. K., Knapp, A. K., … & Gao, W. (2019). Assessing precipitation, evapotranspiration, and NDVI as controls of US Great Plains plant production. Ecosphere, 10(10), e02889.

Cui, T., Martz, L., Lamb, E. G., Zhao, L., & Guo, X. (2019). Comparison of grassland phenology derived from MODIS satellite and PhenoCam near-surface remote sensing in North America. Canadian Journal of Remote Sensing, 45(5), 707–722.

Currier, C. M., & Sala, O. E. (2022). Precipitation versus temperature as phenology controls in drylands. Ecology, 103(11), e3793.

Dai, Y., Zeng, X., Dickinson, R. E., Baker, I., Bonan, G. B., Bosilovich, M. G., … & Yang, Z. L. (2003). The common land model. Bulletin of the American Meteorological Society, 84(8), 1013–1024.

Del Grosso, S. J., Parton, W. J., Keough, C. A., & Reyes-Fox, M. (2011). Special features of the DayCent modeling package and additional procedures for parameterization, calibration, validation, and applications. Methods of introducing system models into agricultural research, 2, 155–176.

Dorich, Christopher D.; Derner, Justin; Torell, Greg; Volesky, Jerry; Brennan, Jameson; Archer, David; Blair, John; Knapp, Alan; Nippert, Jesse; Hartnett, David; McClaran, Mitchel; Maurer, Greg; Moore, Douglas; Clark, Pat; Parton, William; Peck, Dannele; Kramer, Lauren; Smith, William Kolby; Elias, Emile; Fuchs, Brian; Schacht, Walter H.; Hendrickson, John; Harmoney, Keith; Collins, Scott; Baur, Lauren; Porensky, Lauren; Vermeire, Lance; Wilcox, Kevin (2021). Grass-Cast Database - Data on aboveground net primary productivity (ANPP), climate data, NDVI, and cattle weight gain for Western U.S. rangelands. Ag Data Commons. 10.15482/USDA.ADC/1521120

Fawcett, D., Cunliffe, A. M., Sitch, S., O’sullivan, M., Anderson, K., Brazier, R. E., … & Zaehle, S. (2022). Assessing model predictions of carbon dynamics in global drylands. Frontiers in Environmental Science, 10, 790200.

Fox, A. M., Hoar, T. J., Anderson, J. L., Arellano, A. F., Smith, W. K., Litvak, M. E., … & Moore, D. J. (2018). Evaluation of a data assimilation system for land surface models using CLM4. 5. Journal of Advances in Modeling Earth Systems, 10(10), 2471–2494.

Friedl, M., Gray, J., Sulla-Menashe, D. (2022). MODIS/Terra+Aqua Land Cover Dynamics Yearly L3 Global 500m SIN Grid V061 [Data set]. NASA EOSDIS Land Processes Distributed Active Archive Center.

Friedl, M., Sulla-Menashe, D. (2022). MODIS/Terra+Aqua Land Cover Type Yearly L3 Global 500m SIN Grid V061 [Data set]. NASA EOSDIS Land Processes Distributed Active Archive Center.

Gherardi, L. A., & Sala, O. E. (2020). Global patterns and climatic controls of belowground net carbon fixation. Proceedings of the National Academy of Sciences, 117(33), 20038–20043.

Hajek, O. L., & Knapp, A. K. (2022). Shifting seasonal patterns of water availability: ecosystem responses to an unappreciated dimension of climate change. New Phytologist, 233(1), 119–125.

He, Y., Jaiswal, D., Liang, X. Z., Sun, C., & Long, S. P. (2022). Perennial biomass crops on marginal land improve both regional climate and agricultural productivity. GCB Bioenergy, 14(5), 558–571.

Hermance, J. F., Augustine, D. J., & Derner, J. D. (2015). Quantifying characteristic growth dynamics in a semi-arid grassland ecosystem by predicting short-term NDVI phenology from daily rainfall: a simple four parameter coupled-reservoir model. International Journal of Remote Sensing, 36(22), 5637–5663.

Hoover, D. L., Lauenroth, W. K., Milchunas, D. G., Porensky, L. M., Augustine, D. J., & Derner, J. D. (2021). Sensitivity of productivity to precipitation amount and pattern varies by topographic position in a semiarid grassland. Ecosphere, 12(2), e03376.

Hui, D., & Jackson, R. B. (2006). Geographical and interannual variability in biomass partitioning in grassland ecosystems: a synthesis of field data. New phytologist, 169(1), 85–93.

Irisarri, J.G.N., Derner, J.D., Porensky, L.M., Augustine, D.J., Reeves, J. and Mueller, K.E. (2016), Grazing intensity differentially regulates ANPP response to precipitation in North American semiarid grasslands. Ecol Appl, 26: 1370–1380. 10.1890/15-1332

Jones, M. O., Robinson, N. P., Naugle, D. E., Maestas, J. D., Reeves, M. C., Lankston, R. W., & Allred, B. W. (2021). Annual and 16-day rangeland production estimates for the western United States. Rangeland Ecology & Management, 77, 112–117.

Knapp, A. K., Beier, C., Briske, D. D., Classen, A. T., Luo, Y., Reichstein, M., … & Weng, E. (2008). Consequences of more extreme precipitation regimes for terrestrial ecosystems. Bioscience, 58(9), 811–821.

Lauenroth, W.K. and Sala, O.E. (1992), Long-Term Forage Production of North American Shortgrass Steppe. Ecological Applications, 2: 397–403.

Li, X., Melaas, E., Carrillo, C. M., Ault, T., Richardson, A. D., Lawrence, P., … & Young, A. M. (2022). A comparison of land surface phenology in the Northern Hemisphere derived from satellite remote sensing and the Community Land Model. Journal of Hydrometeorology, 23(6), 859–873.

Liang, X. Z., Choi, H. I., Kunkel, K. E., Dai, Y., Joseph, E., Wang, J. X., & Kumar, P. (2005). Surface boundary conditions for mesoscale regional climate models. Earth interactions, 9(18), 1–28.

Liu, Lingling, and Xiaoyang Zhang. "Effects of temperature variability and extremes on spring phenology across the contiguous United States from 1982 to 2016." Scientific Reports 10.1 (2020): 17952.

McCulley, R. L., Burke, I. C., & Lauenroth, W. K. (2009). Conservation of nitrogen increases with precipitation across a major grassland gradient in the Central Great Plains of North America. Oecologia, 159, 571–581.

Milchunas, D. G., & Lauenroth, W. K. (2001). Belowground primary production by carbon isotope decay and long-term root biomass dynamics. Ecosystems, 4, 139–150.

Morgan, J. A., Parton, W., Derner, J. D., Gilmanov, T. G., & Smith, D. P. (2016). Importance of early season conditions and grazing on carbon dioxide fluxes in Colorado shortgrass steppe. Rangeland ecology & management, 69(5), 342–350.

Muldavin, E. H., Moore, D. I., Collins, S. L., Wetherill, K. R., & Lightfoot, D. C. (2008). Aboveground net primary production dynamics in a northern Chihuahuan Desert ecosystem. Oecologia, 155, 123–132.

Nippert, J. B., Ocheltree, T. W., Skibbe, A. M., Kangas, L. C., Ham, J. M., Shonkwiler Arnold, K. B., & Brunsell, N. A. (2011). Linking plant growth responses across topographic gradients in tallgrass prairie. Oecologia, 166, 1131–1142.

NCEP, N. (2005). National Centers for Environmental Prediction (NCEP) North American Regional Reanalysis (NARR). Boulder, CO: Research Data Archive at the National Center for Atmospheric Research, Computational and Information Systems Laboratory

Olson, D. M., Dinerstein, E., Wikramanayake, E. D., Burgess, N. D., Powell, G. V. N., Underwood, E. C., D’Amico, J. A., Itoua, I., Strand, H. E., Morrison, J. C., Loucks, C. J., Allnutt, T. F., Ricketts, T. H., Kura, Y., Lamoreux, J. F., Wettengel, W. W., Hedao, P., Kassem, K. R. (2001). Terrestrial ecoregions of the world: a new map of life on Earth. Bioscience 51(*11*):933*-*938.

Parton, W. J., Schimel, D. S., Cole, C. V., & Ojima, D. S. (1987). Analysis of factors controlling soil organic matter levels in Great Plains grasslands. Soil Science Society of America Journal, 51(5), 1173–1179.

Parton, W. J., Hartman, M., Ojima, D., & Schimel, D. (1998). DAYCENT and its land surface submodel: description and testing. Global and planetary Change, 19(1-4), 35–48.

Paruelo, J.M., Epstein, H.E., Lauenroth, W.K. and Burke, I.C. (1997), ANPP ESTIMATES FROM NDVI FOR THE CENTRAL GRASSLAND REGION OF THE UNITED STATES. Ecology, 78: 953–958.

Petrie, M. D., N. A. Brunsell, R. Vargas, S. L. Collins, L. B. Flanagan, N. P. Hanan, M. E. Litvak, and A. E. Suyker (2016), The sensitivity of carbon exchanges in Great Plains grasslands to precipitation variability, J. Geophys. Res. Biogeosci., 121, 280–294, doi:10.1002/2015JG003205.

Post, A. K., Hufkens, K., & Richardson, A. D. (2022). Predicting spring green-up across diverse North American grasslands. Agricultural and Forest Meteorology, 327, 109204.

Ren, S., Chen, X., Lang, W., & Schwartz, M. D. (2018). Climatic controls of the spatial patterns of vegetation phenology in midlatitude grasslands of the Northern Hemisphere. Journal of Geophysical Research: Biogeosciences, 123(8), 2323–2336.

Reeves, M. C., Hanberry, B. B., Wilmer, H., Kaplan, N. E., & Lauenroth, W. K. (2021). An assessment of production trends on the Great Plains from 1984 to 2017. Rangeland Ecology & Management, 78, 165–179.

Ryals, R., M.D. Hartman, W.J. Parton, M.S. DeLonge, and Silver, and W.L. Silver. (2015). Long-term climate change mitigation potential with organic matter management on grasslands. Ecological Applications, 25(2), pp. 531–545. (10.1890/13-2126.1)

Sala, O. E., Parton, W. J., Joyce, L. A., & Lauenroth, W. K. (1988). Primary production of the central grassland region of the United States. Ecology, 69(1), 40–45.

Schwarz, G. E., & Alexander, R. B. (1995). State soil geographic (STATSGO) data base for the conterminous United States (No. 95-449).

Seyednasrollah, B., Young, A. M., Hufkens, K., Milliman, T., Friedl, M. A., Frolking, S., & Richardson, A. D. (2019). Tracking vegetation phenology across diverse biomes using Version 2.0 of the PhenoCam Dataset. Scientific data, 6(1), 222.

Smith, J. T., Allred, B. W., Boyd, C. S., Davies, K. W., Jones, M. O., Kleinhesselink, A. R., … & Naugle, D. E. (2022). The elevational ascent and spread of exotic annual grass dominance in the Great Basin, USA. Diversity and Distributions, 28(1), 83–96.

Smith, W. K., Dannenberg, M. P., Yan, D., Herrmann, S., Barnes, M. L., Barron-Gafford, G. A., … & Yang, J. (2019). Remote sensing of dryland ecosystem structure and function: Progress, challenges, and opportunities. Remote sensing of environment, 233, 111401.

Sun, Y., Feng, Y., Wang, Y., Zhao, X., Yang, Y., Tang, Z., … & Fang, J. (2021). Field-based estimation of net primary productivity and its above-and belowground partitioning in global grasslands. Journal of Geophysical Research: Biogeosciences, 126(11), e2021JG006472.

White, Robin, Siobhan Murray, and Mark Rohweder, (2000). Pilot Analysis of Global Ecosystems: Grassland Ecosystems, World Resources Institute, Washington D.C. November 2000. ISBN 1-56973-461-5.

Wylie, B., Howard, D., Dahal, D., Gilmanov, T., Ji, L., Zhang, L., & Smith, K. (2016). Grassland and cropland net ecosystem production of the US Great Plains: Regression tree model development and comparative analysis. Remote Sensing, 8(11), 944.

Xia, Y., Sanderman, J., Watts, J., … & Billesbach, D. (2024). Coupling Remote Sensing with a Process Model for the Simulation of Rangeland Carbon Dynamics. ESS Open Archive. (DOI: 10.22541/essoar.171269303.34921116/v1)

