## Supplementary material for "Informing grassland ecosystem modeling with in-situ and remote sensing observations": Supplementary Material.pdf

### SP1 DayCent-UV optimization

The following is a summary of steps we implemented when calibrating DayCent-UV for CPER (short grass steppe) and Konza (tall grass prairie). The parameters calibrated during the optimization process are summarized in Table S1.

- PRDX(1) was most important control on ANPP and was optimized for all our runs. Best fit values range from 0.5 at CPER to 1.0 at Konza.
- EPNFS2(2) was the second most important variable and had a optimum value similar to for both sites (range from 0.6 to 0.8). This controls the amount of non-symbiotic soil N fixation (wdfxs, gN $m^{-2}$ $y^{-1}$) which is calculated annually based on the ratio of actual evapotranspiration (AETan, cm $y^{-1}$) to potential evapotranspiration (PETan, cm $y^{-1}$) wdfxs = EPNFS2(2)*(AETan/PETan) + EPNFS2(1)
- We adjusted the impact of live biomass and soil N levels on C/N ratio of live shoots and roots.
    - PRAMN(1,1)= 20, PRAMN(1,2)=35, PRAMX(1,1)=60, and PRAMX(1,2)=120.
    - PRBMN(1,1)= 60 and PRBMX(1,1)=100 (root C/N ratios).
- We worked on optimizing BIOMAX and ended up with a value of 120 for Konza and 250 for CPER. Higher value for plants with higher N content (short grass vs. tallgrass with many stems).
- We adjusted the impact of water and N stress on carbon allocation to live roots.
    - CFRTCN(1)=0.5, CFRTCN(2)=0.35, CFRTCW(1)=0.75, and CFRTCW(2)=0.45.
- We increased the fraction of live carbon removal from grazing from 0.05 to 0.2 in order to get more removal of live biomass (approximately 50 to 60% of plant production removed by grazers with 0.2/month).
- The amount of C and N returned after fire at Konza was adjusted. 60% of burned N was returned following the fire while 2% of the C was returned. We increased the N return % in order to reduce the negative impact of removing N annually at the tall grass prairie.
- We reduced the number of soil layers where N and water control plant growth (CLAYPG=5 vs. 6 and 7 used earlier). CLAYPG=5, 6, and 7 correspond to a soil depth of 90 cm, 120 cm, and 150 cm, respectively.
- CMIX was increased to 2.5 (mixing of surface slow C into the soil).
- The fraction of N available for plant growth (SFAVAIL(1) ) was increased 0.15 to 0.35 at CPER.
- The soil water controls on plant growth were adjusted to get the impact of drought on plant growth. WSCOEFF(1,1)=0.6 and WSCOEFF(1,2)=9 to 11.
- The impact of water stress on death of live shoots was increased (FSDETH(1)=0.41 for both sites)
- We used the SENM (senescene) schedule file event to get live plant biomass to decrease at the end of the growing season. FSDETH(2) = 0.5 the death rate for each SENM event. SENM events (4 total) were set up to match the observed decrease in live biomass based on the remote sensing NDVI phenology data.

- The model was set up so that the initial soil C equals the values from the Burke equation in the site file (Burke et al. 1989). The fraction of C in the passive carbon was set equal to 0.7 for the Burke equation in the new code.
- We adjusted the root death rates for juvenile and mature live roots. Values for RDRJ are 0.25 for CPER and 0.8 for Konza. RDRM values are 0.06 for CPER and 0.20 for Konza. Higher rates were necessary for Konza in order to get the total live roots correct ( around 750 $gC/m^2$) due to the low water stress at Konza.
- The goal is to have 70% of the soil C in the passive carbon. We will need to make sure that the DEC4 and DEC5(2) values give us the correct ratio. We also need to check the impact of clay content on soil C levels (PS1S3(1) and PS1S3(2)). The flows from slow to passive need to be reduced to low levels based on current knowledge about source of passive C (stabilization of microbe DOC and products on clay particles).
- When simulating the grid, PRDX(1) was linearly increased from the value used for CPER to value used for Konza according to mean annual precipitation (anppt, cm $y^{-1}$) (PRDX(1) = 0.20 + 0.01*anppt).

Table S1. Definitions of DayCent-UV parameters adjusted during calibration.

| **Parameter Name** | **Definition** | **CPER original and calibrated values** | **Konza original and calibrated values** |
|---|---|---|---|
| PRDX(1) | Multiplier to scale radiation use effiency used to calculate potential plant production for crops and grasses. | 0.36 to 0.5 | 2. 0 to 1.0 |
| EPNFS2(2) | Slope term for the effect of the AET/PET ratio on non-symbiotic N fixation. | 0.61 to 0.3 | 0.3 to 0.61 |
| BIOMAX | Aboveground biomass level above which the minimum and maximum C/E ratios of new shoot increments equal PRAMN(*,2) and PRAMX(*,2) respectively. | 200 to 120 | 200 to 250 |
| PRAMN(1,1) | Minimum aboveground C/N ratio of new growth when biomass is zero. | 20 | 20 |
| PRAMN(1,2) | Minimum aboveground C/N ratio of new growth when biomass > BIOMAX | 60 to 35 | 60 to 35 |
| PRAMX(1,1) | Maximum aboveground C/N ratio of new growth when biomass is zero. | 30 to 60 | 40 to 60 |
| PRAMX(1,2) | Maximum aboveground C/N ratio of new growth with biomass > BIOMAX | 80 to 120 | 120 |
| PRBMN(1,1) | (N, intercept) parameter for computing minimum C/N ratio for new growth of belowground matter as a linear function of annual precipitation. (Note: slope was set to 0.0). | 40 to 60 | 60 |
| PRBMX(1,1) | (N, intercept) parameter for computing maximum C/N ratio for new growth of belowground matter as a linear function of annual precipitation (Note: the slope was set to 0.0). | 50 to 100 | 100 |
| CFRTCN(1) | Maximum fraction of C allocated to new root growth under maximum nutrient stress | 0.6 to 0.5 | 0.5 |
| CFRTCN(2) | Minimum fraction of C allocated to new root growth with no nutrient stress | 0.3 to 0.35 | 0.25 to 0.35 |
| CFRTCW(1) | Maximum fraction of C allocated to new root growth under maximum water stress | 0.6 to 0.75 | 0.5 to 0.75 |
| CFRTCW(2) | Minimum fraction of C allocated to new root growth with no water stress | 0.3 to 0.45 | 0.25 to 0.45 |
| CMIX | Annual rate of mixing of surface SOM2C and soil SOM2C ($yr^{-1}$) | 3.0 to 2.25 | 0.25 to 2.25 |
| SFAVAIL(1) | Fraction of inorganic soil N available per day to plants. The remaining fraction is available to microbes. | 0.15 to 0.35 | 0.35 |
| WSCOEFF(1,1) | Water Stress Coefficient used to calculate the water stress multiplier on potential growth based on the relative water content of the wettest soil layer in the rooting zone | 0.6 | 0.45 to 0.6 |

| | | | |
|---|---|---|---|
| WSCOEFF(1,2) | Water Stress Coefficient used to calculate the water stress multiplier on potential growth based on the relative water content of the wettest soil layer in the rooting zone | 25 to 9 | 9 |
| FSDETH(1) | Maximum shoot death rate at very dry soil conditions; to get the monthly shoot death rate, this fraction is multiplied by a reduction factor depending on the soil water status. | 0.04 to 0.4122 | 0.2 to 0.4122 |
| FSDETH(2) | Fraction of shoots which die during each senescence event. | 0.9 to 0.5 | 0.95 to 0.5 |
| RDRJ | Maximum juvenile fine root death rate at very dry soil conditions (fraction/month) | 0.3 to 0.25 | 0.8 |
| RDRM | Maximum mature fine root death rate at very dry soil conditions (fraction/month) | 0.09 to 0.06 | 0.2 |
| DEC4 | Maximum decomposition rate of soil passive organic matter, som3c ($yr^{-1}$). | 0.001 to 0.000597 | 0.001 to 0.000597 |
| DEC5(2) | Maximum decomposition rate of soil slow organic matter; som2c(2) ($yr^{-1}$). | 0.08 to 0.124145 | 0.08 to 0.124145 |

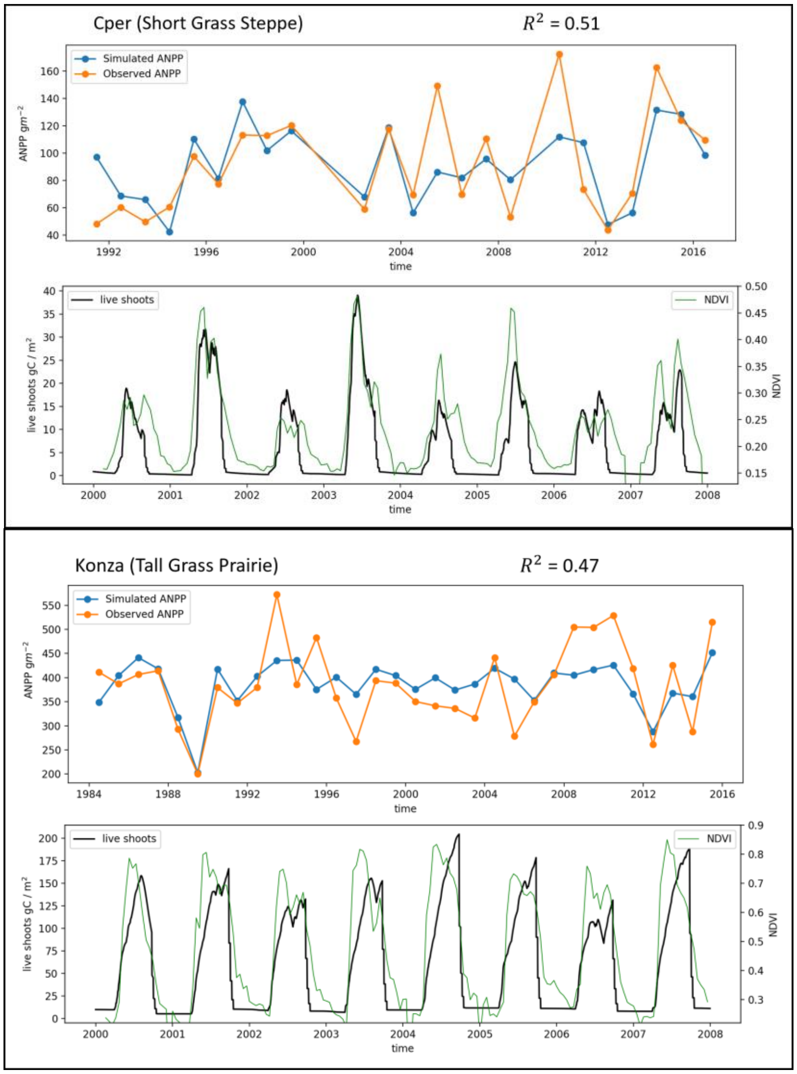


*Figure S1. Final optimization results for CPER and Konza. For each site the top panel represents the temporal ANPP performance of the the simulated ANPP (orange line) respect to site observations (blue line). The bottom panel presents the simulated live shoots production (black line) and the daily MODIS NDVI (green line)*

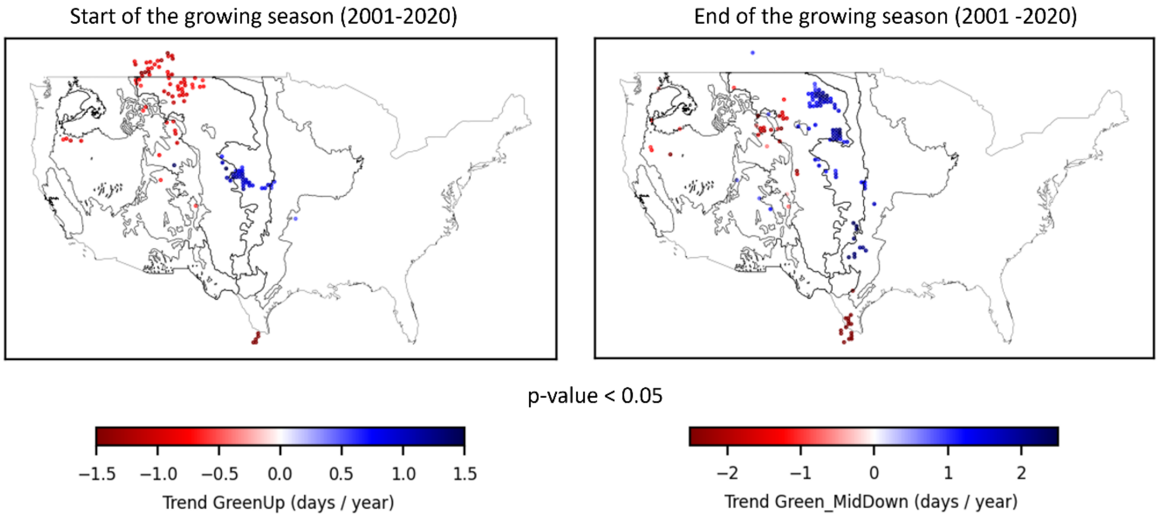


*Figure S2. Significant trends ($p < 0.05$) for the GreenUp band and MidGreenDown band from MODIS MCD12-Q2. Negative slopes (red color) indicates early onset and positive trends (blue color) indicated delayed onset.*

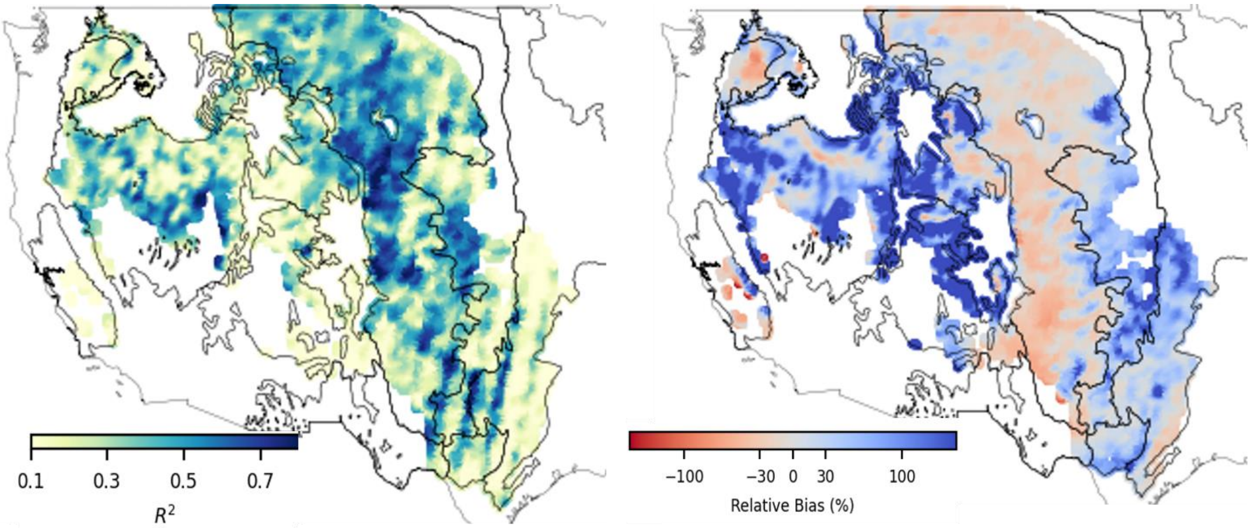


*Figure S3.*

*Temporal $R^2$ between DayCent-UV ANPP and RAP ANPP (left) when the original Eqn 1 was used to estimate RAP ANPP. The corresponding relative bias between DayCent-UV ANPP and RAP ANPP (right)*

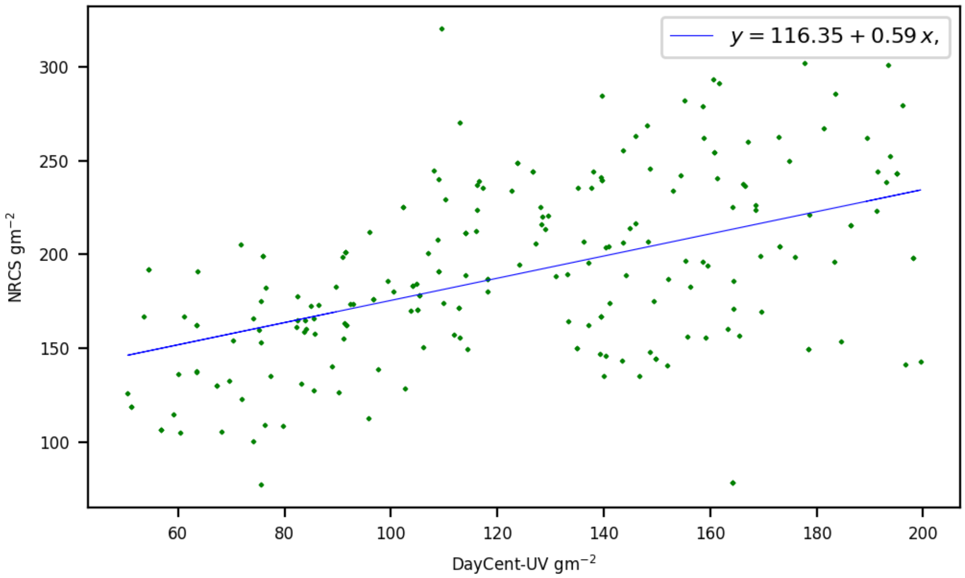


*Figure S4. Bias between NRCS and DayCent-UV at western sites with annual productivity below (200 g/m$^2$)*
